# Association of TSH level above 2.1 mlU/L and first trimester pregnancy loss in anti-TPO antibody negative women

**DOI:** 10.1101/018051

**Authors:** Yisrat Jahan, Enayetur Raheem, Mohammad Akteruzzaman, M Anwar Hussain, Rezaul Karim Kazal, Rifat A. Jahan

## Abstract

Although the exact level of TSH that is indicative of risk of pregnancy loss is not known, a number of studies have suggested a range of values for TSH level that are associated with first trimester pregnancy loss. We conducted an unmatched case-control study to test if a TSH level above 2.1 mlU/L is associated with first trimester pregnancy loss in anti-TPO antibody negative women. We found relatively higher number of women in the case group (18) whose TSH level was above 2.1 mlU/L compared to 7 women in control group. When considered patients in Group I (TSH ≤ 2.1 mlU/L), 45.74% had miscarriage while 54.26% did not have miscarriage within first trimester of pregnancy. Among the Group II patients (TSH > 2.1 mlU/L), 78% had miscarriage and 28% did not have miscarriage. Noticeably there is a larger proportion of miscarriage among the women with TSH level above 2.1 mlU/L. The association between TSH level and first trimester pregnancy loss was statistically significant (p=.0196). From the multivariate analysis, odds ratio for TSH level (OR 4.0, 95% CI: 1.44-11.16) indicates that odds of having miscarriage whose TSH level is above 2.1 mlU/L is 4 times compared to those with TSH level below 2.1 mlU/L after adjusting for the effects of age and BMI. At a global level, the findings of this study provide evidence to the existing discussion on redefining the upper limit of TSH level that is related to first trimester pregnancy loss. At the local level, the results will have direct implication in facilitating management of future pregnancies particularly during the first trimester among Bangladeshi thyroid autoantibody negative women. (268 words)

## INTRODUCTION

The definition of ‘normal TSH level’ during pregnancy is changing. Although TSH values of 0.4 to 5.0 mIU/L were considered normal in the past, studies suggest that first trimester TSH values greater than 2.5 mIU/L, and second and third trimester values greater than 3.00 mIU/L are outside the normal range (Abalovich et al., 2002). The decrease in the upper range of TSH level during first trimester of pregnancy may be attributed to the elevation of human chorionic gonadotropin (hCG), which reacts with the TSH receptor causing a decline in the first trimester TSH level (Hershman, 2004). A recent study found TSH level in the first trimester to be negatively correlated with birth weight of neonates (Kharb, Singla, & Nanda, 2014). This also indicates a potential association between TSH level and its impact on first trimester pregnancy loss.

Although the exact level of TSH that is indicative of risk of pregnancy loss is not known, a number of studies have suggested a range of values for TSH level that are associated with first trimester pregnancy loss. For instance, based on a follow up study involving 343 Chinese women, Panesar et al. (2001) reported a normal range for first-trimester TSH levels of 0.03–2.3 mIU/L. In a study of 585 thyroid antibody-negative women Pearce et al. (2008) found 95% of TSH levels were between 0.04 and 3.6 mIU/L. Study conducted in 1817 Australian women (Gilbert et al., 2008), who were between 9 and 13 weeks of gestation reported a normal TSH range of 0.02–2.15 mIU/L. Stricker et al. (2007) screened 783 thyroid antibody-negative women from the Geneva, Switzerland area and reported a 95% confidence interval for TSH level to be 0.08–2.83 mIU/L. Although the reported ranges of TSH level to be considered as ‘normal’ vary, there are consistencies among the study results. There is no agreed upon value of the TSH level, however, a consensus on a lower limit of normal to be 0.04 and upper limit of normal being 2.5 (Negro et al., 2010).

It is important to consider clinical implications such as pregnancy loss and preterm delivery of untreated cases with first trimester TSH in a range that had previously been considered normal. The increase incidence of pregnancy loss in pregnant women with TSH level between 2.0 and 5.0 mIU/L provides justification to consider investigating the TSH upper limit of normal in the first trimester to a value around 2.0 mIU/L. For the purpose of present study, we chose to evaluate the impact of TSH level above 2.1 mIU/L (high normal) on first trimester pregnancy loss. Of particular interest is to study the socioeconomic determinants that might be associated with first trimester pregnancy loss along with TSH level. Our study involved socioeconomically mid- to well-off women in Dhaka city in Bangladesh. We hypothesize that a TSH value above 2.1 mlU/L may be associated with first trimester pregnancy loss in anti TPO Ab negative women.

## MATERIALS AND METHODS

To test our research hypothesis, a cross sectional case control study was conducted where the subjects were selected from out-patient clinics of three centers in Dhaka, Bangladesh. The centers are Department of Obstetrics and Gynaecology under Bangabandhu Sheikh Mujib Medical University (BSMMU), Dhaka Medical College Hospital (DMCH), and Mohammadpur Fertility Services and Training Centre (MFSTC). After screening for the inclusion and exclusion criteria patients were recruited for the study as they arrived at the facility. Informed written consent was taken from each participant. A predesigned data collection sheet was used to collect data. Ethical clearance for this study was taken from the Institutional Review Board (IRB) of BSMMU.

### Inclusion criteria for cases and controls

The case group contained Bangladeshi born anti-TPO Ab negative women between 18-45 years of age, gestational period up to first trimester (12 weeks), and who had recent pregnancy loss which was diagnosed clinically and confirmed by USG. Women in the control group had the similar inclusion criteria but with continuing pregnancy. Anti-TPO Ab level < 35 IU/mL was considered as negative antibody.

### Exclusion criteria

Women with the following conditions were not included in the study: pregnancy with known thyroid disease; pregnant women suffering from acute illness such as viral hepatitis, typhoid; those suffering from chronic diseases such as chronic hypertension, chronic renal disease, uncontrolled diabetes mellitus; women who were on medication that interfere with the thyroid function such as steroid, carbamezapine. Further, women who lost pregnancy due to other causes were excluded from the study.

Patients were assigned to case group if there was miscarriage during first trimester. Otherwise, they were assigned to the control group. Subjects were further categorized into two groups according to their TSH level. Group I with TSH level at or below 2.1 mlU/L and those with TSH level above 2.1 mlU/L were in Group II.

### Data and variables

After selection of the study subjects, history taking and clinical examination were performed, and diagnosis was confirmed by USG. Medical history, clinical examination results as well as demographic and socioeconomic variables such as patients’ age, height, weight, family income, and husband’s occupation were collected.

To determine TSH level, 5 ml of venous blood was drawn from ante-cubital vein using disposable syringe with all aseptic precaution. Blood samples were transferred immediately into clean and dry test tubes with gentle push after removal of the needle to avoid hemolysis. The samples were allowed to clot and then centrifuged. Serum was aliquoted into label micro-centrifuged tubes and preserved at 2-8 degree Celsius for future analysis. Analyses of TPO-Ab and TSH were completed using Abott Axsym system/AxSYM 3^rd^ Generation TSH assay within five days of sampling.

### Statistical Analysis

Bivariate and multivariate analyses were carried out to the study the association between TSH level and first trimester pregnancy loss. The descriptive results are reported as mean ± SD. Student’s ‘t’-test was performed to test for differences in the means, and Chi-square test was used to test for association between variables. Bivariate results are shown in Tables 1 and 2.

**Table 1:** Cross-tabulation of TSH level and exposure to event (miscarriage).

|  | **Case**<br>**n (%)** | **Control**<br>**n (%)** | **Chi-square**<br>**(p-value)** |
| --- | --- | --- | --- |
| Group I<br>TSH ≤ 2.1 mIU/L | 43<br>(45.74)<br>(70.49) | 51<br>(54.26)<br>(87.93) | 5.45<br>(.0196) |
| Group II<br>TSH $>$ 2.1 mIU/L | 18<br>(72.00)<br>(29.51) | 7<br>(28.00)<br>(12.07) | |

Figures in each cell represent the counts (first row), row percentage (second row), and column percentage (third row).

**Table 2:** Cross-tabulation family income and exposure to event (miscarriage).

| **Family Income** | **Case**<br>**n (%)** | **Control**<br>**n (%)** | **Chi-square**<br>**(p-value)** |
| --- | --- | --- | --- |
| $<$ 70,000 Taka/year | 27<br>(57.14)<br>(44.26) | 36<br>(42.86)<br>(62.07) | 3.78<br>(.0517) |
| ≥ 70,000 Taka/year | 34<br>(60.71)<br>(55.74) | 22<br>(39.29)<br>(37.93) | |

Figures in each cell represent the counts (first row), row percentage (second row), and column percentage (third row).

Multivariate analysis was performed to obtain adjusted odds ratios using a binary logistic regression model. For the multivariate analysis, age of patients, BMI, TSH category (≤ 2.1 mlU/L vs > 2.1 mlU/L), and family income (<70,000 Taka/year vs ≥ 70,000 Taka/year) were considered as covariates. Maximum likelihood estimates of the effects and adjusted odds ratios are presented in Table 3.

**Table 3:** Estimated effects, adjusted odds ratios and 95% confidence intervals for the odds ratios.

| Maximum Likelihood Estimates |  |  |  |  | Odds Ratio Estimates |  |  |
| --- | --- | --- | --- | --- | --- | --- | --- |
| Parameter | Estimate | Standard Error | Wald Chi-Square | Pr > ChiSq | Point Estimate | 95% Wald Confidence Limits |  |
| Intercept | -0.21 | 1.24 | 0.03 | 0.8685 |  |  |  |
| Age | -0.02 | 0.04 | 0.18 | 0.6688 | 0.98 | 0.91 | 1.06 |
| BMI | -0.002 | 0.04 | 0.004 | 0.9478 | 0.99 | 0.93 | 1.07 |
| TSH level (>2.1) | 1.39 | 0.52 | 7.05 | 0.0079 | 4.01 | 1.44 | 11.16 |
| Family Income<br>(≥ 70,000 Taka/yr) | 0.97 | 0.41 | 5.70 | 0.0169 | 2.64 | 1.19 | 5.85 |

The results in this paper were generated using SAS/STAT software, Version 9.4 of the SAS System for Windows. Copyright, SAS Institute Inc. SAS and all other SAS Institute Inc. product or service names are registered trademarks or trademarks of SAS Institute Inc., Cary, NC, USA.

## DISCUSSION

In this study, we had 58 controls and 61 cases. Patients who had a miscarriage within the first trimester were in case group, and who did not have any miscarriage during the first trimester were in the control group. Patients were carefully selected so that the case and control groups are comparable. However, the subjects in case and control groups were not matched. On the basis of serum TSH level, the subjects were further classified as ‘low normal’ (TSH level up to 2.1 mIU/L) and ‘high normal’ (TSH level above 2.1 mIU/L). The case and control groups were similar with respect to background characteristics as shown in Table 1. Average ± SD for age (in years) of patients in the case group was 25.06 ± 5.36, and for the control group it was 24.68 ± 5.07. Negro et al. (2010) in their study, however, had relatively higher mean age both in the cases and controls (28.7 and 29.2 respectively). Average weight (kg) for case group was 55.45 ± 7.58 and for control group 55.13 ± 6.95. Mean BMI for case group was 25.24 ± 6.13, and for the control group it was 24.98 ± 4.31. The two groups were comparable with respect to gestational age (weeks); 8.46 ± 1.88 for the case group and 9.32 ± 2.27 for the control group. We found no statistically significant difference between case and control groups for any of the above mentioned indicators.

The study found the number of women to be relatively higher in Group II (TSH level > 2.1 mIU/L) for the cases (18) compared to the controls (7). Among the cases, 29.51% had TSH level above 2.1 mlU/L whereas 12.07% of the control group patients had TSH level above 2.1 mlU/L. When considered patients in Group I (TSH ≤ 2.1 mlU/L), 45.74% had miscarriage while 54.26% did not have miscarriage. Within the Group II patients, 78% had miscarriage and 28% did not have miscarriage. Noticeably there is a larger proportion of miscarriage that had TSH level above 2.1 mlU/L. Indeed, the association between TSH level and exposure to event (miscarriage) was statistically significant (Chi-square=5.45, p=.0196).

Cross tabulation of family income (below 70,000 Taka/year vs above 70,000 Taka/year) by event status is shown in Table 2. In the case group, 55.74% had family income above 70,000 Taka per year while the remaining 44.26% had income less than 70,000 Taka per year. Overall, there is no noticeable difference between the case and control groups based on the income categories. We found marginal association (p=.0517) between family income and exposure to miscarriage which warrants for further investigation through multivariate analysis.

Since univariate or bivariate results do not take into consideration the effects of other factors that might affect the event of interest (miscarriage), we fit a multiple logistic regression model considering age, BMI, family income, and TSH level as independent variables. The outcome variable is binary with two categories indicating whether miscarriage occurred or not. The results are presented in Table 3. We found family income and TSH level to be significant predictors of miscarriage. However, age and BMI were not significant. Adjusted odds ratio for TSH level (OR 4.0, 95% CI: 1.44-11.16) indicates that odds of having miscarriage whose TSH level is above 2.1 mlU/L is 4 times compared to those with TSH level below 2.1 mlU/L after adjusting for the effects of age and BMI. The 95% confidence interval confirms the significance of the estimated odds ratios. Further, adjusted odds ratio for family income (OR 2.64, 95% CI: 1.19-5.85) indicates that adjusted odds of having miscarriage whose family income above 70,000 Taka/year is 2.64 times compared to those with lower family income.

## CONCLUSION

It was well documented in the reviewed literature that TSH level has significant effect on the first trimester pregnancy loss among the thyroid autoantibody negative women. However, the upper limit varies across different demographic characteristics of women. Proper maternal thyroid function during pregnancy is important for both the mother and the developing fetus. This is particularly true during the first trimester, when the fetus is completely dependent on the mother for thyroid hormone. This study detects significant relationship between high normal TSH (>2.1 mlU/L) and miscarriage in first trimester among mid- to well-off women whose family income is above 70,000 Taka/year. At a global level, our findings provide evidence to the existing discussion on redefining the upper limit of TSH level that is related to first trimester pregnancy loss. At the local level, the study results will have direct implication in facilitating management of future pregnancies particularly during the first trimester among Bangladeshi thyroid autoantibody negative women.

## ACKNOWLEDGEMENTS

The authors thank the anonymous referees for their valuable comments which have improved the presentation of the paper.

